# Abiotic treatment to common bean plants results in an altered endophytic seed microbiome

**DOI:** 10.1101/2020.06.05.134445

**Authors:** A. Fina Bintarti, Patrick J. Kearns, Abby Sulesky-Grieb, Ashley Shade

**Affiliations:** Department of Plant, Soil and Microbial Sciences, Michigan State University, East Lansing MI 48824; The Plant Resilience Institute, Michigan State University, East Lansing MI 48824; Department of Microbiology and Molecular Genetics, Michigan State University, East Lansing MI 48824

**Keywords:** plant microbiome, 16S rRNAgene, ITS, drought, fertilizer, legume, growth chamber, endophyte, vertical transmission, abiotic stress

## Abstract

There has been a growing interest in the seed microbiome due to its important role as an end and starting point of plant microbiome assembly that can have consequences for plant health. However, the effect of abiotic conditions on the seed microbial community remains unknown. We performed a pilot study in a controlled growth chamber to investigate how the endophytic seed microbiome of the common bean *(Phaseolus vulgaris L.* (var. Red Hawk)) was altered under abiotic treatments relevant for crop management with changing climate. Bean plants were subjected to one of three treatments: 66% water withholding to simulate mild drought, 50% Hoagland nutrient solution to simulate fertilization, or control with sufficient water and baseline nutrition. We performed 16S rRNA gene amplicon sequencing and ITS1 amplicon sequencing of the endophytic DNA to assess seed bacterial/archaeal and fungal community structure, respectively. We found that variability in the seed microbiome structure was high while alpha diversity was low, with tens of taxa present. Water withholding and nutrient addition altered the seed microbiome structure for bacterial/archaeal communities as compared to the control, and each treatment resulted in a distinct microbiome structure. There were no statistically supported differences in the fungal microbiome across treatments. While we discuss several limitations of this study, the promising results suggest that further investigation is needed to better understand abiotic or stress-induced changes in the seed microbiome, the mechanisms that drive those changes, and their implications for the health and stress responses of the next plant generation.

**Importance:** Seed microbiome members initiate the assembly of plant-associated microbial communities, but the environmental drivers of endophytic seed microbiome composition are unclear. Here, we exposed plants to short-term drought and fertilizer treatments during early vegetative growth and quantified the microbiome composition of the seeds that were ultimately produced. We found that seeds produced by plants stressed by water limitation or receiving nutrient addition had statistically different endophytic bacterial/archaeal microbiome compositions from each other and from seeds produced by control plants. This work suggests that the abiotic experience of a parental plant can influence the composition of its seed microbiome, with unknown consequences for the next plant generation.

## Introduction

The plant microbiome includes bacteria, archaea, fungi, and viruses that associate with the plant and inhabit different plant compartments, including the rhizosphere, phyllosphere, and endosphere (1). The plant microbiome plays important roles for plant fitness, including nutrient acquisition (2), secondary metabolite production (3), flowering time (4), and resistance to abiotic (5, 6) and biotic stresses (7, 8). Plant microbiota can interact with each other as well as with the host plant, and the plant is able to shape its microbiome structure and composition, for example, by producing root exudates or allelochemicals (9, 10). Because of the close relationship between plants and their associated microbiomes, it is likely that both or either could be affected by external stressors and, in turn, also affect each other. These environmental stressors can include abiotic stress, such as changes in water and nutrient availability (limitation or excess) or exposure to extreme temperatures, and biotic stress, such as pathogen infection and herbivory. As plants respond to various stressors, their microbiome may also be altered, either as a direct or indirect consequence of the stress. A recent study reported increases in stress-related gene expression in the rhizosphere microbial community of plants treated with high pH and high salinity wastewater, which suggested that the functional gene profile and expression pattern of the plant microbiome under stressors can be used as an indicator tool to identify stresses affecting host plants (11). Thus, both the environmental conditions and the host plant can act as important filters that contribute to the ultimate composition of the plant microbiome (12, 13).

As a critical part of the plant microbiome, the seed microbiome can directly impact the seed and seedling in ways that are important for crop establishment, such as by releasing the seed from dormancy and promoting seed germination and seedling emergence (14,15). However, there are relatively fewer comprehensive studies of the seed microbiome as compared to studies of the rhizosphere and phyllosphere microbiomes, which can acquire microbiota from the environment, (e.g., aerosols (12) and soils (16, 17)). Previously, investigation of seed microbiota employed culture-dependent methods and focused on the transmission of plant pathogenic bacteria or fungi (18, 19). More recently, seed microbiome studies have expanded and adopted cultivation-independent methods (20–25). It is now recognized that the seed represents an endpoint of microbiome assembly for the parental plant’s reproductive compartment and the starting point of microbiome assembly for the new seedling (26).

Vertical transmission of microbes via the seed has been reported for a variety of plant species, as recently summarized (26) and reported (23, 27). Seeds acquire microbiota through different modes of transmission, where early colonizers of the seed endophytic environment are acquired from the parental plant either through the vascular system or floral stigma, while late colonizers are acquired on external surfaces via seed contact with the environment (24, 28). A recent study of temporal dynamics of the bean seed microbiome assembly reported that the vascular pathway is the dominant route for seed microbiome transmission in common bean (24). Seed endophytes, acquired through the parental plant vascular tissue, are of great interest because they are vertically transmitted to plant offspring, and plants may preserve specific taxa through vertical transmission over generations. We hypothesized that these preserved taxa may play vital roles in plant growth and tolerance to environmental stress. Thus, in this study, we focused on seed endophytes and excluded seed epiphytes.

Managing or manipulating the plant microbiome is one promising strategy to support plant tolerance to environmental stress. We are just beginning to understand how the plant microbiome structure is altered during particular stresses (e.g., drought (29)), with a focus in the literature on the root zone and rhizosphere microbiome. An important initial step of plant microbiome engineering to enhance plant fitness and growth under environmental stresses is to understand the effect of these stresses on the plant microbiome, and, next, to decipher underlying mechanisms involved in the process. A previous study revealed that grass root microbiome diversity and structure was affected by drought and there was enrichment for Actinobacteria (30). Drought resulted in reduced diversity of sorghum root microbiome and increased abundance and activity of Gram-positive, and this shift was correlated with altered plant metabolism and increased expression of bacterial ATP-binding cassette transporter genes (29). Another recent study showed shifts in the wheat seed microbiome where Actinobacteria were enriched and Gammaproteobacteria were depleted under drought conditions, and these selected seed microbiome members demonstrated plant growth promoting ability on plants undergoing drought (31). These studies that we highlight here are a few of many papers in this area of the root and rhizosphere microbiome and its response to stress, but they do not investigate the seed microbiome.

We conducted a controlled pilot study in an environmental growth chamber to determine changes in the rhizosphere microbiome of the legume common bean under two different treatments of water withholding and nutrient addition. The initial purpose of our pilot study was to identify members of the root microbiome that were particularly resilient to either of these treatments. However, the plants set pods at the end of the experiment, and we realized the opportunity to also assess the seed microbiome of the treated plants as compared to control plants. The purpose of this brief report is to share the surprising seed microbiome results from the pilot study, to discuss its limitations, and to suggest immediate future directions based on the most promising results.

## Methods

### Plant growth conditions and harvest

Common bean seeds of the Red Hawk cultivar (32) were obtained from a laboratory in the Michigan State University Plant Biology department. Seeds were surface-sterilized in a 10% bleach solution followed by five rinses in sterile DI water. Three seeds per pot were planted in 24 one-gallon pots filled with a steam-sterilized (~100°C) mixture of agricultural topsoil, sphagnum peat and sand, and culled to one seedling per pot after the first unifoliate leaves had emerged. The plants were grown in controlled conditions in a high-light BioChambers FLEX™ LED growth chamber with a 16-hour day/8-hour night cycle at 26°C and 22°C, respectively. The plants were divided into three groups: 8 control plants received ample water (300 mL every other day), 8 plants were subjected to a mild “drought” during plant development, and received 66% less water (100 mL every other day) (water withholding), and 8 plants received half strength Hoagland solution (300 mL every other day) provided by the growth chamber facility (nutrient addition). Hoagland Solution details can be found in Supplementary Table 1. The plants were grown for ~60 days until the R7 stage, when plant pods were fully developed. All plants grew at relatively the same rate as expected for the cultivar, so were harvested at the same time.

Harvesting was conducted by collecting the bean pods and plant biomass. Bean pods were removed from the plants and the remaining above ground biomass from each plant was placed in a paper bag and dried at 70°C for one week. The root system was gently pulled from the pot, cleaned of excess soil with deionized water and dried at 70°C for one week. Once dried, the shoot and root dry weight was measured for each plant. The remaining soil was collected for soil chemical analysis. One hundred grams of each soil sample along with three replicates of pre-treatment bulk soil were sent to the Michigan State University Soil and Plant Nutrient Laboratory (SPNL) for soil chemical testing. Soil parameters including pH, lime index, phosphorus (P), potassium (K), calcium (Ca), magnesium (Mg), nitrate (NO_3_^-^), ammonium (NH_4_^+^), and organic matter (OM) were measured for all soil samples (testing procedures available in Supplementary Methods, data available in Supplementary Table S2). Soil chemistry differences among treatments were assessed using one-way analysis of variance (ANOVA). The normality and homoscedasticity of the data were evaluated using Shapiro-Wilk and Levene’s test, respectively. We performed non-parametric Kruskal-Wallis test and post-hoc Dunn’s test with Benjamini-Hochberg correction for *p*-values if the ANOVA’s assumptions were not met.

### DNA extraction and amplicon sequencing

Twenty seeds from each plant were collected for DNA extraction following the protocol of a previous study (20) with minor modifications to include surface sterilization (33). Seeds were surfaced sterilized in 10% bleach, rinsed five times with sterile DI water, and placed in sterile 50 mL centrifuge tubes with 30 mL of sterile 1X phosphate buffered saline (PBS) with 0.05% Tween 20 and shaken at 140 rpm at room temperature for 4 hours. After shaking, tubes were centrifuged at 500 x g for 15 minutes and the supernatant and seeds were discarded. The remaining pellet was resuspended with 2 mL of sterile 1X PBS-Tween and transferred to a microcentrifuge tube and spun at 20,000 x g for 10 min. The supernatant was discarded, and the pellet was used for DNA extraction with the PowerSoil□DNA Isolation Kit (MoBio Laboratories, Solana Beach, CA, United States) following manufacturer’s instructions. DNA extracted from seed samples was quantified with Qubit™dsDNA BR Assay Kit (ThermoFisher Scientific, Waltham, MA, United States) and verified with Polymerase Chain Reaction (PCR), including a negative PCR control and blank extraction reagents.

The bacterial/archaeal community PCR was conducted using the 515f (5’-GTGCCAGCMGCCGCGGTAA-3’) and 806r (5’-GGACTACHVGGGTWTCTAAT-3’) primer pair (34) for amplification of the V4 region of the 16S rRNA gene. The 16S rRNA gene amplification was conducted under the following conditions: 94°C for 3 min, followed by 34 cycles of 94°C (45 s), 50°C (60 s), and 72°C (90 s), with a final extension at 72°C (5 min). The amplification was performed in 25 μl mixtures containing 12.5 μl GoTaq□Green Master Mix (Promega, Madison, WI, United States), 0.625 μl of each primer (20mM), 1 μl of DNA template (final concentration of 0.02 – 0.626 ng per μl), and 4.5 μl nuclease free water. Seed DNA (concentration range of 5 – 20 ng per μl) was sequenced at the Research Technology Support Facility (RTSF) Genomics Core, Michigan State University sequencing facility using the Illumina MiSeq platform.

Fungal communities were assessed using PCR amplification of the Internal Transcribed Spacer 1 (ITS1) region with the ITS1□ (5□-CTTGGTCATTTAGAGGAAGTAA-3□) and ITS2 (5□-GCTGCGTTCTTCATCGATGC-3□) primer pair (35) with the addition of index adapters as required by the RTSF Genomics Core, (https://rtsf.natsci.msu.edu/genomics/sample-requirements/illumina-sequencing-sample-requirements/). The PCR conditions of the ITS gene amplification conditions were as follows: 95°C for 5 min, followed by 30 cycles of 95°C (30 s), 54°C (45 s), and 72°C (90 s), with a final extension at 72°C (5 min). The amplification was performed in 50 μl mixtures containing 20 μl GoTaq□Green Master Mix (Promega, Madison, WI, United States), 1 μl of each primer (20 mM), 4 μl of DNA template (final concentration of 0.02 – 0.626 ng per μl), and 26 μl nuclease free water. The product of the ITS gene amplification was cleaned and purified using the Wizard□SV Gel and PCR Clean-Up System (Promega, Madison, WI, United States), following the manufacturer’s protocol. Purified ITS gene amplification products with the concentration range of 5 – 50 ng per μl were sequenced at the RTSF Genomics Core using the llumina MiSeq platform. No amplification of the ITS gene was observed in one water withholding and one nutrient addition seed sample, so only seven samples were sequenced for fungal analysis in these treatment groups.

The 16S and ITS libraries were prepared by the sequencing facility using the Illumina TruSeq≡Nano DNA Library Prep Kit (Illumina, Inc., San Diego, CA, United States), llumina MiSeq was run using a v2 Standard sequencing format with paired end reads (2 x 250 bp), and negative and mock positive sequencing controls provided by the sequencing facility were included with each run.

### Sequencing data analysis and OTU clustering

Bacterial/archaeal raw reads produced from Illumina MiSeq were processed including merging the paired end reads, filtering the low-quality sequences, dereplication to find unique sequence, singleton removal, denoising, and chimera checking using the USEARCH pipeline (v.10.0.240) (36). Operational taxonomic unit (OTU) clustering was conducted using an open reference strategy (37). First, closed reference OTU picking was performed at 97 % identity by clustering quality filtered reads against the SILVA database (v.132) (38) using USEARCH algorithm (-*usearch_global* command) (39). Reads that failed to match the SILVA reference were subsequently clustered *de novo* at 97% identity using UPARSE-OTU algorithm *(-cluster_otus* command) (40). Closed reference and *de novo* OTUs were combined into a full set of representative sequences, and then all merged sequences were mapped back to that set using the *-usearch_global* command.

The set of representative sequences were aligned on QIIME 1.9.1 (41) using PyNAST (42) against the SILVA (v.132) reference database. The unaligned OTU sequences were excluded from the OTU table and the representative sequences. Taxonomic assignment was conducted on QIIME 1.9.1 using the SILVA (v.132) database and the UCLUST default classifier at a minimum confidence of 0.9 (39). Plant contaminants such as chloroplast and mitochondria; and unassigned taxa and sequences were removed from the OTU table as well as the representative sequences *using filter_taxa_from_otu_table.py* and *filter_fosta.py* command on QIIME. Rarefaction to the lowest sequencing depth (43, 44) (11,137 bacterial/archaeal reads) was conducted on QIIME.

The processing of fungal ITS raw reads was also conducted using the USEARCH (v.10.0.240) pipeline. Read processing included paired end read merging, primer removal using cutadapt (v.2.0) (45), filtering the low-quality sequences, and dereplication to find unique sequence. Operational taxonomic unit clustering was conducted using an open reference OTU picking strategy. First, closed reference OTU picking was performed by clustering quality filtered reads against the UNITE fungal ITS database (v.8.0) (46) at 97% identity threshold using the USEARCH algorithm. Reads that failed to match the reference were clustered *de novo* at 97% identity using the UPARSE-OTU algorithm. Closed reference and *de novo* OTUs were combined into a full set of representative sequences, and then all merged sequences were mapped back to that set using *-useorch_globol* command. Fungal taxonomic classification was performed in the CONSTAX tool (47) at a minimum confidence of 0.8 using the UNITE reference database release 01-12-2017. Assigning taxonomy in CONSTAX was conducted using three classifiers, including RDP Classifier (v.11.5) (48, 49), UTAX from USEARCH (v.8.1.1831) (40), and SINTAX from USEARCH (v.9.2) (50). Any contaminants including mitochondria, chloroplast and other unwanted lineages of eukaryotes were removed from the OTU table. Rarefaction was conducted to the lowest number of sequences (21,329 fungal reads) on QIIME.

### Microbial community analysis

Microbial community analyses were conducted in R (v.3.6.1) (R Core Development Team). Microbial composition and relative abundance were analyzed using the Phyloseq package (v.1.28.0) on R (51). Microbial richness (the number of taxa present) was calculated on the rarefied OTU table using the vegan package (v.2.5-6) (52). The normality and homoscedasticity of the data were tested using Saphiro-Wilk and Levene’s test, respectively. The one-way ANOVA or non-parametric Kruskal-Wallis test was then performed to analyze the data. Post hoc Dunn’s test with false discovery rate (FDR) correction using the Benjamini-Hochberg adjustment for multiple comparisons was performed to compare plant biomass data among treatments.

Beta diversity was calculated on the rarefied OTU table with the vegan package using Jaccard dissimilarity indices and visualized with a principal coordinate analysis (PCoA) plot. We used the Jaccard index, which is based on presence-absence counts rather than relative abundance data, because we reasoned that the seed microbiome members are unlikely to be actively growing inside the seed and that any differences in relative abundances in the seed endophyte are unlikely attributable to competitive growth outcomes *in situ.* Permutational multivariate analysis of variance (PERMANOVA) using the function adonis (52) was performed to assess the effects of the treatments to the microbial community structure. We performed multivariate analysis to check the homogeneity of dispersion (variance) among groups using the function betadisper (52).

We analyzed shared microbial taxa between seeds from control and treated plants by calculating the their occupancy (53). Microbial OTUs with occupancy value of 1 were those OTUs that were detected in all samples from included treatments (control and nutrient addition, control and water withholding).

### Data and code availability

The computational workflows for sequence processing and ecological statistics are available on GitHub (https://github.com/ShadeLab/BioRxiv_Seed_Microbiome_2020). Raw sequence data of bacteria/archaea and fungi have been deposited in the Sequence Read Archive (SRA) NCBI database under Bioproject accession number PRJNA635871.

## Results

There were overall differences in plant biomass among treatments (Fig. 1, detailed Kruskal-Wallis results can be found in Supplementary Table S3). Specifically, plants receiving nutrient addition were larger in shoot and root biomass than control or mildly droughted plants (Fig. 1). Nutrient addition plants also had higher pod number and pod mass compared to the water withholding and control plants (Fig. 1). As expected, the addition of the Hoagland solution increased the nutrients available to the plants in the nutrient addition treatment as compared to the control, as rhizosphere soil from the nutrient addition treatment had higher phosphorus and potassium content than the other two treatments,as well as higher nitrate content than the control treatment (Supplementary Figure S1, Supplementary Table S4).

**Fig 1.**
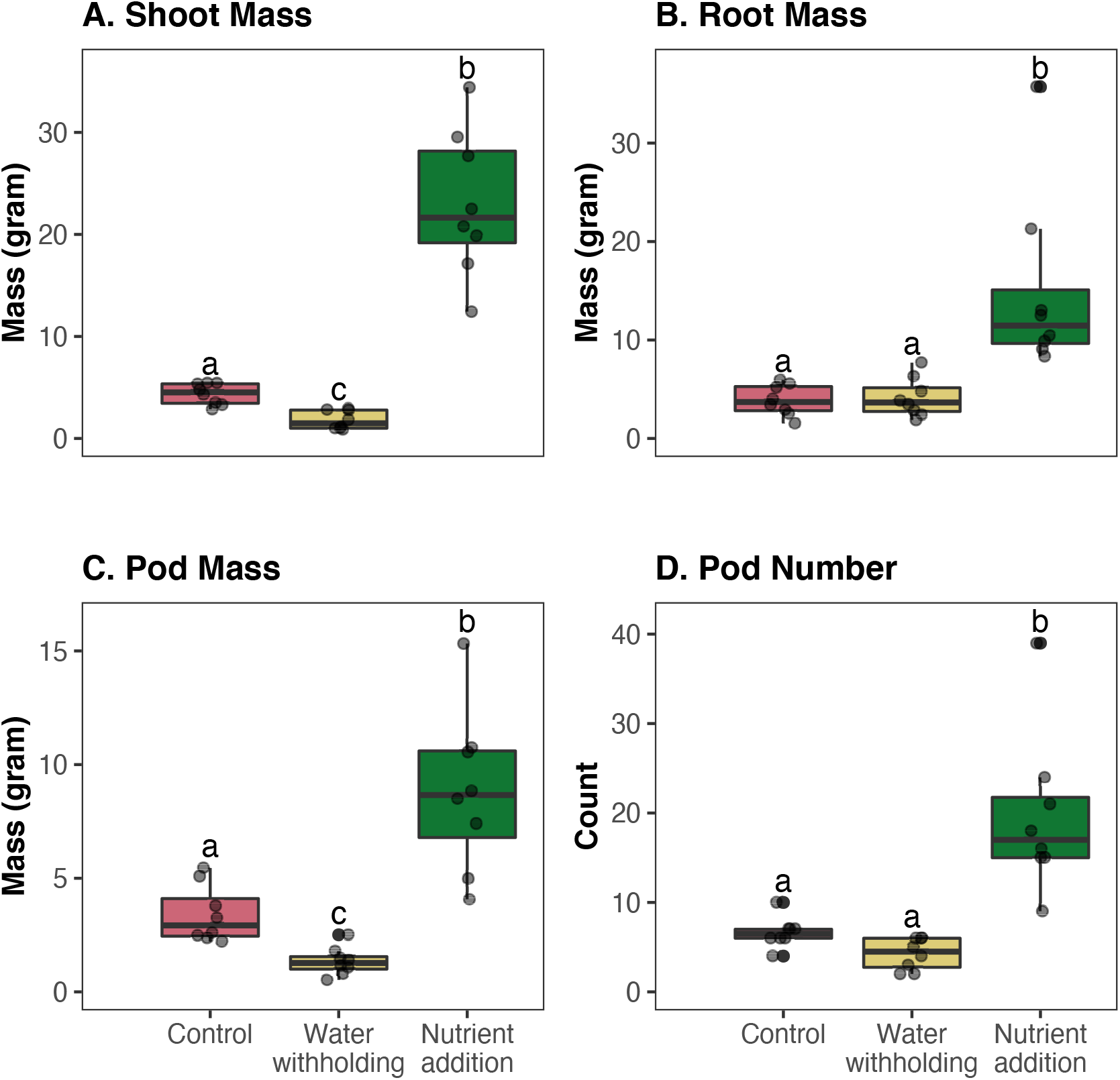
Plant aboveground (shoot) and belowground (root) biomass for control, water withholding, and nutrient addition treatments of common bean. Plant biomasses were calculated on eight plant replicates for each treatment. For each box plot, circles represent a single plant measurement within a treatment. The central horizontal lines represent the mean, the outer horizontal lines of the box represent the 25th and 75th percentiles. Boxes labelled with different letters were significantly different by a Kruskal-Wallis and post-hoc Dunn’s test with a Benjamini-Hochberg false discovery rate correction (p-value significance ranges from <0.05 to <0.0001).

We removed less than 0.1 % of reads identified as plant and eukaryote contaminants from bacterial/archaeal and fungal sequences, respectively. Analysis of contaminant-filtered bacterial/archaeal and fungal sequences from seed samples resulted in a total of 81 and 226 OTUs (97% sequence identity), respectively. Bacterial/archaeal communities in control, water withholding, and nutrient addition seeds had different taxonomic compositions (Fig. 2A). Bacterial/archaeal communities in the control seeds were almost exclusively dominated by the OTUs within the genus *Bacillus,* with a mean relative genus-level abundance of more than 99%. Although the bacterial/archaeal community in the water withholding and nutrient addition seeds were also dominated by *Bacillus,* genus-level taxonomic diversity increased with the addition of other, non-dominating lineages. Specifically, seed communities from water withholding and nutrient addition plants were also composed of *Virgibacillus, Pseudomonas,* and several other bacterial/archaeal genera.

**Fig 2.**
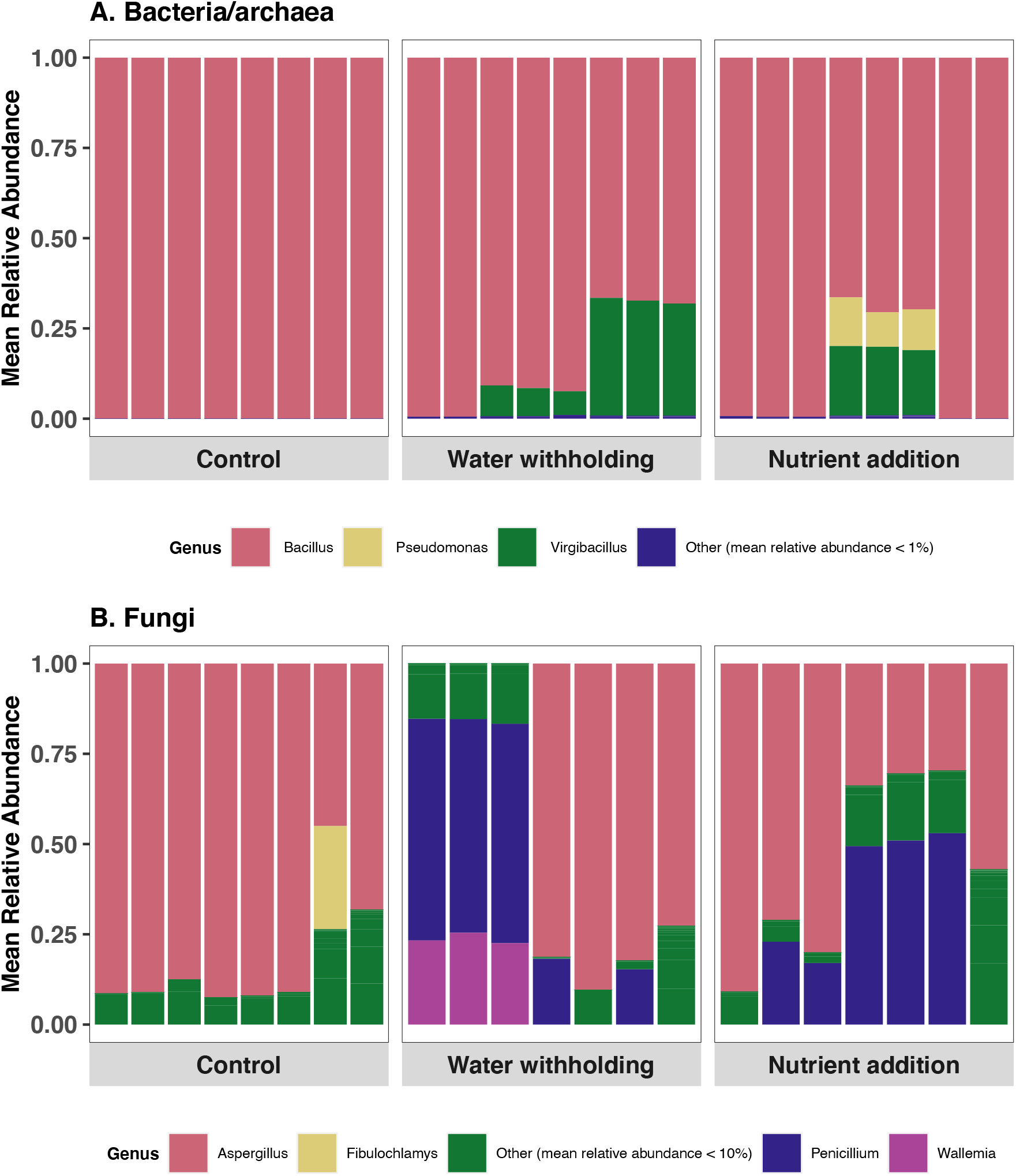
Mean relative abundances of genera of bacteria/archaea (A) and fungi (B) detected in the seed across control, water withholding and nutrient addition treatments. Each bar represents the endophytic microbiome identified in DNA extracted from 20 seeds collected from one plant replicate within a treatment. Bacterial/archaeal and fungal genera with mean relative abundances of less than 1 and 10 %, respectively, were grouped into the ‘Other’ classification, which includes many lineages (not monophyletic). Genera identified in the ‘Other’ classifications can be found in Supplementary Tables S5 and S6.

Similarly, different plant treatments resulted in different seed fungal community compositions. Even though *Aspergillus* dominated the fungal community in the control and treated seeds, in the treated seeds there was a shift to include other fungal taxa, including some identified as *Penicillium* and *Wallemia* (Fig. 2B). These observations indicate that the seed microbiome is altered when parental plants are exposed to abiotic stress or environmental alteration.

Analysis of overlapping taxa across treatments revealed that there were 4 and 3 bacterial taxa shared between control and water withholding-treated seeds, and between control and nutrient addition seeds, respectively (Supplementary Table S7). Bacterial taxa shared between samples belonged exclusively to genus *Bacillus.* Fungal communities were dominated by genus *Aspergillus* that were detected between control and water withholding-treated seeds, and *Penicillium* and *Aspergillus* that were shared between control and nutrient addition seeds (Supplementary Table S8).

Because we do not expect microbiome members to be actively doubling inside the seed (and therefore relativized abundances inside the seed to reflect fitness differences as an outcome of growth therein, see (33) or a discussion of this), we used a presence-absence assessment (Jaccard index) of beta diversity. There was a statistically supported differences in bacterial/archaeal microbiome composition between treated seeds and control seeds (Fig. 3A, permutational multivariate analysis of variance, PERMANOVA, F-stat = 4.73, R^2^ = 0.31, P-val = 0.001). In contrast, there was no distinct clustering of fungal communities associated with different treatments (Fig. 3B, PERMANOVA P-val > 0.05). These results indicate that the abiotic treatments applied in this study altered the bacterial/archaeal, but not fungal, community composition in the common bean seed. Also, there were differences in the composition variability (multivariate dispersion) among treatments for bacterial/archaeal communities (permutational analysis of multivariate dispersion, PERMDISP, F=7.553, P-val=0.003). Specifically, the seeds from plants that experienced nutrient addition and water withholding had higher dispersion as compared to seeds from control plants (TukeyHSD.betadisper, P-val=0.003 and 0.03, respectively). Meanwhile, there were no differences in dispersion observed for fungal composition (PERMDISP, F=0.491, P-val=0.628). This provides additional evidence that these abiotic treatments can lead to increased variability in seed microbiome composition. Notably, PERMANOVA was found to be largely unaffected by heterogeneity for balanced designs (54).

**Fig. 3.**
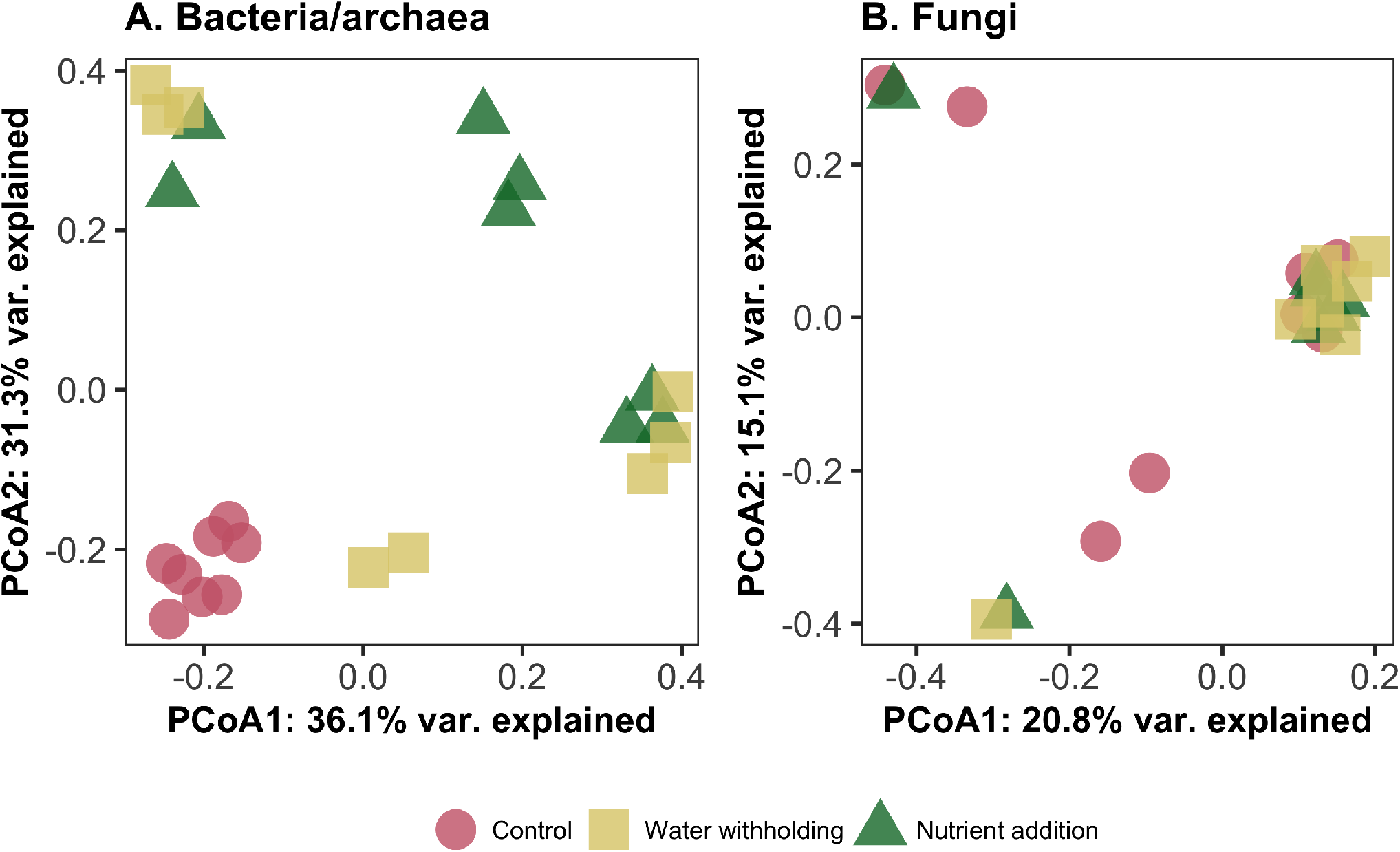
Principal coordinate analysis (PCoA) plot of the bacterial/archaeal community (PERMANOVA, F-stat = 4.73, R^2^ = 0.31, P-val = 0.001) (A) and fungal community (PERMANOVA P-val > 0.05) (B) in the common bean seed based on the Jaccard index. Symbol colors and shapes represent different abiotic treatments on the parent plant.

## Discussion

The identities, functions and persistence of seed microbiome members are either not known or not well-understood for many plant species of ecological or agricultural importance. This fundamental information is the first step in understanding what constitutes a “typical” microbiome to initiate plant microbiome assembly from seed. Changes to a typical seed microbiome because of parental plant exposure to stress may have consequences for the health or resilience of the next plant generation. For example, a depletion of beneficial members or enrichment of pathogens in the seed could disadvantage the plant offspring, while an enrichment of beneficial members could provide a health advantage. Therefore, observing a modification in the seed microbiome after parental exposure to stress has potentially important implications and warrants report and continuation of research to understand any consequences in functions for plant health.

We highlight three important observations from this study. First, treatment of the parent plant altered the seed endophyte structure and composition compared to control plants, especially for the bacterial/archaeal community. Although studies on the impact of drought on seed microbiota are still new, especially as compared to studies on drought and the root-associated microbiome, our results are consistent with a recent work which reported shifts in seed microbial communities of wheat under drought conditions (31). Shifts in microbial communities have also been observed in various studies on the root and rhizosphere of different plant species under drought stress (6, 55–57) with some reporting that the effect of drought was stronger on the endophyte communities relative to rhizosphere communities (30, 55, 57). This suggests an indirect effect of drought to endophyte communities through physiological changes in drought-affected plants.

To our knowledge, this study is the first study assessing the impact of nutrient addition on seed endophytes, as previous studies into the effect of fertilization have focused almost exclusively on the rhizosphere and root-associated microbiome. We observed alterations in the seed endophyte communities from nutrient addition parental plants. A previous study on the common bean rhizosphere microbiome reported higher diversity in agricultural soil than native soil, suggesting that management practices including fertilization may be a driver of the observed differences (58). Additionally, our recent work on the biogeography of common bean rhizosphere microbiomes across different bean production regions in the U.S revealed that fertilization differences between samples (including synthetic, organic manure and no-input) had an explanatory value on the microbial community structure (59).Together, these results suggest that any perturbations that affect the host plant may also affect its microbial communities and these alterations may result in specific outcomes for plant stress tolerance and resilience. Since the seed microbiomes contained a relatively simple community of tens to dozens of taxa, even alterations in the composition or abundances of a few taxa may have consequences for microbiome assembly of the next plant generation.

Our second key observation in this study is that bacterial/archaeal communities from the seeds of treated plants had more variation as compared to the seeds from control plants. This suggests that abiotic stress results in changes analogous to those observed during other types of microbiome “dysbiosis” (aka Anna Karenina effects: higher variability across replicates, increased beta-dispersion, and higher contribution of stochastic assembly processes (60)). As observed in dysbiosis of human or animal microbiomes, the plant microbiome structure can be disrupted by disturbances and changing environmental conditions that alter the composition and diversity of ‘normal’ microbiota (61, 62).

However, unlike dysbiosis in the human microbiome, any deviations from unaltered microbial communities are not always associated with negative effects on plant health, and this alteration may be necessary for maintaining plant health under stress conditions (63). Notably, there is no clear definition of a “healthy” plant microbiome, though it has been suggested that high diversity and high evenness are typical characteristics of healthy plant microbiomes and dysbiosis is correlated with reduced diversity (64–66). However, we know that diversity metrics are aggregations of complex community data that are intended to compare but ultimately directly cannot explain underlying biological mechanisms (67). We speculate that the variations observed in the seed endophyte composition in our study, and the increase in alpha and beta diversity of wheat seed microbiome of drought-exposed plants from another recent study (31), are possibly due to selection of functionally beneficial microbial taxa. It has been hypothesized that the selection and enrichment of particular taxa under drought stress is more likely associated with functionality rather than taxonomy (56). There could be a microbial shift towards a state associated with positive effects on plant health, as opposed dysbiosis (63)). However, the expectation of high variability should be taken into consideration for future studies, as sufficient replication will be needed to power statistical tests (e.g., (33)).

Third and finally, the fungal community was, on balance, stable relative to the bacterial/archaeal community, suggesting that the persistence of fungal members is less sensitive to water withholding and nutrient addition treatments. It has been reported that fungi can be very resistant to environmental disturbances including drought due to several mechanisms including thick cell wall and osmolytes (68–70). A previous study reported that drought had a more pronounced impact on bacteria than on fungal networks in soil (71). Another study also reported that bacterial communities were significantly and consistently altered not only in composition and diversity but also co-occurrence networks in forests disturbed by clear-cutting or conversion to agricultural use relative to non-disturbed, meanwhile these effects were marginal in fungal communities (72). All together, this suggests that fungal communities are more stable over bacterial community under environmental changes.

We acknowledge that this is a pilot study and that these results are preliminary. We offer a discussion of some of the major considerations and limitations in interpreting the results and for planning future seed microbiome studies.

A first consideration is that there is an apparent maximum stress to plants that can be applied when investigating its consequence fora seed microbiome. After stress exposure is released, plants must be healthy enough to produce pods and seeds, and a balance must be achieved in which plants are stressed but still able to fully mature. This constraint in stress exposure will never accommodate an experimental design of severe or prolonged stress. However, the investigation of a mild or moderate stress is still valuable because it is pertinent to agriculture. There are many situations in which non-lethal stress occurs over part of a growing season, but then crops recover fully or partially to produce some yield. Therefore, the result of mild or moderate stress for seed microbiomes has real-world relevance.

Another consideration is the definition and directness of abiotic treatment, and whether an abiotic treatment is expected to act on the plant, the microbiome, or both. In this pilot study, we applied two different abiotic treatments: one that was expected to stress the plant directly (water withholding to simulate mild drought) and one that was expected to weaken a legume’s relationship with its root-associated microbiome and symbiotic nitrogen fixers as nitrogen fixers are down-regulated by nitrogen application (73, 74) (nutrient addition). Thus, the addition of nutrients was a benefit to the plant, rather than a stress, as indicated by the increased root and shoot biomass. However, nutrient addition caused a clear shift in the seed microbiome, demonstrating the potential of fertilizer use to have multi-generational impacts on plant microbiome assembly. Therefore, management practices that advantage the plant as far as yield and health in the short-term could have long-term consequences for plant-microbiome relationships.

A clear limitation of the study is the substrate used for plant growth, which, with the microbes in and on the original seeds, serves as a starting source for the assembly of the new plant’s microbiome (20, 26). For this pilot study, we used a sterilized mixture of agricultural topsoil, sphagnum peat and sand provided by the growth chamber facility. The exact origin and physical/chemical characteristics of the facility soil is unknown, and so it is unclear how representative this soil may be of agricultural field soil. Additionally, the initial microbial community in the soil was not analyzed before planting the common bean plants, so we cannot determine the origin of the observed microbial consortia in the seeds and to what extent they overlap with the potting substrate. Previous work suggests that soil type can have a large influence on the seed endophytic bacteria in rice (75), and this is likely also true for other plant seeds. We observed a dominance of taxa from genus *Bacillus* in common bean seeds in all three treatments, and *Bacillus* taxa have previously been reported to be enriched in the green bean seedling (20). However, steam sterilization of the growth chamber soil may have killed many indigenous microbial taxa in the soil that were not spore-formers or otherwise resistant to heat (and, bacilli are known to be resistant to such treatments, as per (76)). Therefore, the microbial consortia available to colonize the plant and seed microbiome from the sterilized soil was likely more limited as compared to plants grown in the field or other substrates. We urge caution in generalizing from specific compositional changes, but rather focus on the larger changes in beta-diversity and dispersion that were consistently observed across very different abiotic treatments and may be more characteristic of seed microbiome responses. Future work should focus efforts on using soil that is representative of the typical agricultural environment of common bean, and the existing microbial community in the soil should be sequenced prior to planting for comparison to the seed microbiome.

*We* included many standard controls in this study, including DNA extraction, PCR, and other standard molecular biology controls. However, another limitation of this study the absence of negative controls for DNA extraction that were followed the whole way through to sequencing. Seed endophytes contain a very low total biomass of microbial cells. Here, we pooled twenty seeds to use for one extraction to increase the microbial biomass yield for microbiome interrogation. We performed a buffer-only control that was PCR-negative, but we did not save the material for sequencing, which would allow for direct assessment of contaminants from the DNA extraction process. While the surface-sterilization of the seeds prior to extraction and negative PCR controls provide confidence that the starting material was not compromised and that we did not unintentionally amplify contaminants from the PCR reagents, we cannot know if there were a few OTUs signatures from the extraction kit or buffer contaminants that could have contributed to the observed seed microbiome composition (though we did not observe any common contaminants reported in the literature). We now advocate for sequencing the DNA extraction buffer control and using a package such as decontam (77) and microDecon (78) to ensure removal of spurious contaminants, which are expected in low biomass samples (79).

A final minor limitation is in the choice of bacterial marker gene. We performed amplicon sequencing of the 16S rRNA gene for the bacterial/archaeal community analysis in the seed. It is well known that the variability of rRNA copy numbers among bacterial species can lead to an inflation of species richness and obfuscate relative abundances of taxa (80). Moreover, the most resolved taxonomic level achieved from 16S rRNA amplicon gene sequences often is genus, rather than species or strain. Using a single-copy marker gene that has higher precision and sensitivity at the species level, like the *gyrB* gene, may be a valuable alternative (20). The *gyrB* gene has been successfully applied to other seed microbiome studies (20, 21, 81, 82). It may be valuable to consider use of both marker genes to the same seed microbiome samples, so that taxonomic precision can be maximized and compared across seeds with *gyrB,* while also maintaining an ability to source-track and compare composition across the many plant-microbiome 16S rRNA amplicon datasets that have been deposited publicly.

In summary, while this pilot study provides a key insight into the response of the seed microbiome structure to abiotic treatment in the host plant, there is much more work to be done. Next steps include exposing the plants to more severe drought and nutrient excess conditions, quantifying the physiological status of plants to determine their experience of stress, using representative field soil for plant growth and assessing the field soil microbiome to deduce seed taxon origins, sequencing negative controls from the DNA extractions to identify contaminants, and considering use of an alternative marker gene for improved precision in microbial taxonomy and taxon abundances.

Despite noted considerations and limitations, we posit that this pilot study revealed an important insight regarding how seed microbiomes may be altered after abiotic treatment of a plant. Next, we need to understand the implications of this change for both the host plant and the microbial community. An altered seed microbiome may have positive, negative, or entirely neutral outcomes for the next plant generation. Additional work is needed to understand these outcomes over consecutive plant generations to determine the effects on plant fitness and resilience. If positive or negative outcomes are detected, this work opens a new direction of research that could spur exciting applications in plant microbiome management.

## Acknowledgements

This work was supported by the Michigan State University Plant Resilience Institute and by the USDA National Institute of Food and Agriculture award USDA 2019-67019-29305. AFB acknowledges support from the Fulbright Foreign Student Program and Office for International Students and Scholars (OISS), Michigan State University; and AS acknowledges support from the USDA National Institute of Food and Agriculture and Michigan AgBioResearch (Hatch). We thank Matthieu Barret and the Shade Lab for helpful discussions.

