## Supplementary for "Abiotic treatment to common bean plants results in an altered endophytic seed microbiome"

by

A. Fina Bintarti, Patrick J. Kearns, Abby Sulesky-Grieb, and Ashley Shade

### Supplementary Tables

**Table S1: One-half strength Hoagland Solution recipe provided by the Michigan State University growth chamber facility.**

| Nutrient | Grams of nutrient per 40L solution |
|---|---|
| **Solution A*** | |
| Calcium nitrate | 11800.00 |
| Sequestrene | 1537.60 |
| **Solution B*** | |
| Potassium phosphate, monobasic | 1370.00 |
| Potassium nitrate | 5066.00 |
| Magnesium sulphate | 2500.00 |
| Zinc sulphate | 2.24 |
| Manganous sulphate | 15.64 |
| Copper sulphate | 0.84 |
| Boric acid | 29.00 |
| Molybdic acid | 0.20 |
| *Both solution A and B are used at 1:500 ratios (76.9 ppm) | |
| KOH is added to adjust the final nutrient solution pH | |

**Table S2: Soil pH, organic matter content (OM) and nutrient content for untreated bulk soil and post-treatment soil collected after plant harvest.**

| Sample | Treatment | pH | P (ppm) | K (ppm) | Ca (ppm) | Mg (ppm) | OM (%) | NO3 (ppm) | NH4 (ppm) |
|---|---|---|---|---|---|---|---|---|---|
| **bulk 1** | none | 7.3 | 32 | 39 | 1506 | 154 | 3.1 | 25.6 | 7.1 |
| **bulk 2** | none | 7.3 | 35 | 39 | 1627 | 170 | 3.2 | 25.9 | 7.6 |
| **bulk 3** | none | 7.3 | 38 | 39 | 1673 | 174 | 3.1 | 24.6 | 6.4 |
| **n7** | nitrogen | 7.7 | 36 | 94 | 1659 | 187 | 3.4 | 28.3 | 0.7 |
| **n6** | nitrogen | 7.4 | 73 | 96 | 1532 | 161 | 3.8 | 24.8 | 0.7 |
| **n5** | nitrogen | 7.6 | 86 | 153 | 1587 | 162 | 4.8 | 21.8 | 0.4 |
| **n4** | nitrogen | 7.4 | 69 | 97 | 1527 | 157 | 3.4 | 26.7 | 0.9 |
| **n3** | nitrogen | 7.6 | 69 | 141 | 1533 | 149 | 2.9 | 19.0 | 1.4 |
| **n2** | nitrogen | 7.6 | 63 | 162 | 1509 | 156 | 3.2 | 29.6 | 1.5 |
| **n1** | nitrogen | 7.7 | 46 | 104 | 1651 | 201 | 3.9 | 20.7 | 1.7 |
| **n8** | nitrogen | 7.7 | 42 | 46 | 1679 | 178 | 3.2 | 11.5 | 0.1 |
| **d1** | drought | 7.9 | 32 | 39 | 1638 | 172 | 3.1 | 3.0 | 0.7 |
| **d2** | drought | 7.7 | 34 | 41 | 1644 | 184 | 3.3 | 37.3 | 1.4 |
| **d3** | drought | 7.7 | 31 | 39 | 1599 | 176 | 4.2 | 2.9 | 1.3 |
| **d4** | drought | 7.8 | 26 | 39 | 1547 | 160 | 3.2 | 3.8 | 0.8 |
| **d5** | drought | 7.6 | 28 | 39 | 1636 | 176 | 3.4 | 2.5 | 0.8 |
| **d6** | drought | 7.8 | 31 | 39 | 1649 | 189 | 3.3 | 21.2 | 1.1 |
| **d7** | drought | 7.7 | 30 | 39 | 1752 | 187 | 4.1 | 3.4 | 1.0 |
| **d8** | drought | 7.8 | 34 | 43 | 1588 | 176 | 3.0 | 42.4 | 1.4 |
| **c1** | control | 7.7 | 29 | 39 | 1648 | 158 | 3.2 | 0.6 | 1.4 |
| **c2** | control | 8.0 | 30 | 39 | 1607 | 164 | 3.1 | 6.7 | 1.2 |
| **c3** | control | 7.9 | 34 | 38 | 1657 | 135 | 3.4 | 0.5 | 1.0 |
| **c4** | control | 7.9 | 27 | 38 | 1574 | 129 | 3.3 | 0.5 | 1.2 |
| **c5** | control | 7.9 | 32 | 38 | 1699 | 163 | 3.7 | 0.9 | 0.7 |
| **c6** | control | 7.9 | 32 | 38 | 1567 | 151 | 3.6 | 0.3 | 0.6 |
| **c7** | control | 7.8 | 30 | 38 | 1669 | 157 | 3.6 | 0.8 | 1.4 |
| **c8** | control | 7.9 | 26 | 39 | 1747 | 219 | 4.9 | 2.4 | 1.1 |

**Table S3: Comparison of plant biomass among treatments. Bolded rows are significantly different as determined by Kruskal-Wallis test and post-hoc Dunn's test with Benjamini-Hochberg false discovery rate (FDR) correction.**

| Kruskal-Wallis test | | | | Post-hoc Dunn's test with Benjamini-Hochberg FDR correction | | | |
|---|---|---|---|---|---|---|---|
| **Parameter** | **Chi-squared** | **df** | **p-val** | **Comparison** | **Z** | **P. unadj** | **P. adj** |
| **Shoot mass** | 20.165 | 2 | **0.00004** | Control – Nutrient addition | -2.29810 | 0.02156 | **0.03233** |
| | | | | Control – Water withholding | 2.19203 | 0.02838 | **0.02838** |
| | | | | Nutrient addition – Water withholding | 4.49013 | 0.00001 | **0.00002** |
| **Root mass** | 15.365 | 2 | **0.00046** | Control – Nutrient addition | -3.42947 | 0.00060 | **0.00181** |
| | | | | Control – Water withholding | -0.07071 | 0.94363 | 0.94363 |
| | | | | Nutrient addition – Water withholding | 3.35876 | 0.00078 | **0.00117** |
| **Pod mass** | 18.305 | 2 | **0.00010** | Control – Nutrient addition | -2.08597 | 0.03698 | **0.03698** |
| | | | | Control –Water withholding | 2.19203 | 0.02838 | **0.04257** |
| | | | | Nutrient addition – Water withholding | 4.27800 | 0.00002 | **0.00006** |
| **Pod number** | 17.973 | 2 | **0.00013** | Control – Nutrient addition | -2.53347 | 0.01129 | **0.01694** |
| | | | | Control – Water withholding | 1.67708 | 0.09353 | 0.09353 |
| | | | | Nutrient addition – Water withholding | 4.21055 | 0.00003 | **0.00008** |

**Table S4: Comparison of rhizosphere soil chemistry of the three treatments using one-way ANOVA or Kruskal-Wallis test. Bolded rows are significantly different as determined by Kruskal-Wallis test and post-hoc Dunn's test with Benjamini-Hochberg false discovery rate (FDR) correction. Post-hoc tests were not performed for samples with insignificant p-values from Kruskal-Wallis or ANOVA tests.**

| | Kruskal-Wallis test | | | Post-hoc Dunn's test with Benjamini-Hochberg FDR correction | | | |
|---|---|---|---|---|---|---|---|
| **Parameter** | **Chi-squared** | **df** | **p-value** | **Comparison** | **Z** | **P. unadj** | **P. adj** |
| pH | 14.634 | 2 | **0.00066** | Control – Nutrient addition | 3.824263 | 0.00013 | **0.00039** |
| | | | | Control – Water withholding | 1.83057 | 0.06716 | 0.06716 |
| | | | | Nutrient addition – Water withholding | -1.99369 | 0.04619 | 0.06928 |
| P | 15.613 | 2 | **0.00041** | Control – Nutrient addition | -3.60033 | 0.00032 | **0.00095** |
| | | | | Control – Water withholding | -0.39018 | 0.69640 | 0.69640 |
| | | | | Nutrient addition – Water withholding | 3.21014 | 0.00133 | **0.00199** |
| K | 19.172 | 2 | **0.00007** | Control – Nutrient addition | -4.34149 | 0.00001 | **0.00004** |
| | | | | Control –Water withholding | -1.67822 | 0.09330 | 0.09330 |
| | | | | Nutrient addition – Water withholding | 2.66327 | 0.00774 | **0.01161** |
| $NO3^-$ | 14.261 | 2 | **0.0008** | Control – Nutrient addition | -3.67776 | 0.00024 | **0.00071** |
| | | | | Control –Water withholding | -2.58150 | 0.00984 | **0.01476** |
| | | | | Nutrient addition – Water withholding | 1.09625 | 0.27297 | 0.27297 |
| Mg | 4.8514 | 2 | 0.08842 | | | | |
| OM | 0.53985 | 2 | 0.7634 | | | | |
| | **One-way ANOVA test** | | | | | | |
| **Parameter** | **F-value** | **df** | **p-value** | | | | |
| Ca | 2.035 | 2 | 0.156 | | | | |
| $NH4^+$ | 0.343 | 2 | 0.713 | | | | |

**Table S5: Bacterial genera identified in <1% of the OTUs in each seed microbiome sample.**

| Treatment | Sample | Bacterial Genus |
|---|---|---|
| **Control** | C1 | Unidentified Steroidobacteraceae |
| | C2 | Rhizobiales |
| | C4 | Solibacillus |
| | C5 | Virgibacillus |
| | | Brevibacillus |
| | | Sporosarcina |
| | C7 | Brevibacillus |
| | C8 | Brevibacillus |
| **Water withholding** | D1 | Dyadobacter |
| | | Anaerobacillus |
| | | Virgibacillus |
| | | Paenibacillus |
| | | Solibacillus |
| | D2 | Anaerobacillus |
| | | Solibacillus |
| | D3 | Anaerobacillus |
| | | Bacillaceae |
| | | Solibacillus |
| | | Sporosarcina |
| | D4 | Anaerobacillus |
| | | Bacillaceae |
| | | Solibacillus |
| | D5 | Anaerobacillus |
| | | Bacillaceae |
| | | Solibacillus |
| | D6 | Pedobacter |
| | | Anaerobacillus |
| | | Bacillaceae |
| | | Sporosarcina |
| | | Escherichia-Shigella |
| | D7 | Dyadobacter |
| | | Anaerobacillus |
| | | Bacillaceae |
| | | Sporosarcina |
| | | Escherichia-Shigella |
| | D8 | Anaerobacillus |
| | | Bacillaceae |
| | | Domibacillus |
| | | Solibacillus |
| | | Sporosarcina |
| | | Escherichia-Shigella |
| | | Pseudomonas |
| **Nutrient addition** | N1 | Blastococcus |
| | | Promicromonospora |
| | | Anaerobacillus |
| | | Solibacillus |
| | N2 | Pedobacter |
| | | Anaerobacillus |
| | | Solibacillus |
| | N3 | Anaerobacillus |
| | | Solibacillus |
| | | Nitrospira |
| | N4 | Sphingobacterium |
| | | Anaerobacillus |
| | | Bacillaceae |
| | | Fictibacillus |
| | | Paenibacillus |
| | | Sporosarcina |
| | | Colwelliaceae |
| | | Escherichia-Shigella |
| | N5 | Sphingobacterium |
| | | Anaerobacillus |
| | | Bacillaceae |
| | | Fictibacillus |
| | | Paenibacillus |
| | | Solibacillus |
| | | Sporosarcina |
| | | Colwelliaceae |
| | | Escherichia-Shigella |
| | N6 | Sphingobacterium |
| | | Anaerobacillus |
| | | Bacillaceae |
| | | Fictibacillus |
| | | Paenibacillus |
| | | Sporosarcina |
| | | Colwelliaceae |
| | | Escherichia-Shigella |
| | N7 | Blastococcus |
| | | Bacillaceae |
| | | Virgibacillus |
| | N8 | Virgibacillus |
| | | Sporosarcina |

**Table S6: Fungal genera identified in <10% of the OTUs in each seed microbiome sample.**
**OTUs that could not be identified at the Genus level are identified at the Order or Family level as labeled below.**

| Treatment | Sample | Fungal Genus |
|---|---|---|
| **Control** | C1 | Alternaria |
| | | Order: Pleosporales |
| | | Penicillium |
| | | Family: Aspergillaceae |
| | | Family: Nectriaceae |
| | | Order: Hypocreales |
| | | Conocybe |
| | | Fibulochlamys |
| | | Ischnoderma |
| | | Filobasidium |
| | | Mortierella |
| | | Unidentified |
| | C2 | Order: Pleosporales |
| | | Penicillium |
| | | Family: Aspergillaceae |
| | | Peziza |
| | | Meyerozyma |
| | | Verticillium |
| | | Family: Nectriaceae |
| | | Order: Hypocreales |
| | | Conocybe |
| | | Fibulochlamys |
| | | Parasola |
| | | Guehomyces |
| | | Filobasidium |
| | | Wallemia |
| | | Mortierella |
| | | Unidentified |
| | C3 | Penicillium |
| | | Family: Aspergillaceae |
| | | Family: Chaetomiaceae |
| | | Sistotrema |
| | | Guehomyces |
| | | Wallemia |
| | | Unidentified |
| | | Unidentified |
| | C4 | Unidentified |
| | | Penicillium |
| | | Family: Aspergillaceae |
| | | Family: Nectriaceae |
| | | Order: Hypocreales |
| | | Apodus |
| | | Wallemia |
| | | Unidentified |
| | C5 | Alternaria |
| | | Order: Pleosporales |
| | | Penicillium |
| | | Family: Aspergillaceae |
| | | Family: Trichocomaceae |
| | | Meyerozyma |
| | | Fusarium |
| | | Gibberella |
| | | Family: Nectriaceae |
| **Nutrient Addition** | N1 | Alternaria |
| | | Order: Pleosporales |
| | | Unidentified |
| | | Penicillium |
| | | Family: Aspergillaceae |
| | | Family: Pyronemataceae |
| | | Family: Nectriaceae |
| | | Order: Hypocreales |
| | | Unidentified |
| | | Marasmius |
| | | Wallemia |
| | | Olpidium |
| | N2 | Alternaria |
| | | Order: Pleosporales |
| | | Unidentified |
| | | Family: Aspergillaceae |
| | | Emericellopsis |
| | | Gibberella |
| | | Family: Nectriaceae |
| | | Order: Hypocreales |
| | | Unidentified |
| | | Fibulochlamys |
| | | Coprinellus |
| | | Guehomyces |
| | | Wallemia |
| | | Mortierella |
| | N3 | Alternaria |
| | | Order: Pleosporales |
| | | Unidentified |
| | | Family: Aspergillaceae |
| | | Verticillium |
| | | Gibberella |
| | | Family: Nectriaceae |
| | | Order: Hypocreales |
| | | Unidentified |
| | | Guehomyces |
| | | Wallemia |
| | N4 | Alternaria |
| | | Order: Pleosporales |
| | | Unidentified |
| | | Family: Sclerotiniaceae |
| | | Family: Pyronemataceae |
| | | Cyberlindnera |
| | | Fusarium |
| | | Gibberella |
| | | Family: Nectriaceae |
| | | Order: Hypocreales |
| | | Order: Microascales |
| | | Humicola |
| | | Apodus |
| | | Family: Lasiosphaeriaceae |
| | | Unidentified |
| | | Parasola |
| **Water Withholding** | D1 | Alternaria |
| | | Drechslera |
| | | Order: Pleosporales |
| | | Unidentified |
| | | Aspergillus |
| | | Family: Aspergillaceae |
| | | Family: Pyronemataceae |
| | | Verticillium |
| | | Fusarium |
| | | Gibberella |
| | | Family: Nectriaceae |
| | | Order: Hypocreales |
| | | Unidentified |
| | | Conocybe |
| | | Fibulochlamys |
| | | Cyathus |
| | | Order: Auriculariales |
| | | Sistotrema |
| | | Uncobasidium |
| | | Udeniozyma |
| | | Guehomyces |
| | | Unidentified |
| | | Mortierella |
| | | Family: Mortierellaceae |
| | | Olpidium |
| | D2 | Alternaria |
| | | Order: Pleosporales |
| | | Unidentified |
| | | Order: Chaetothyriales |
| | | Aspergillus |
| | | Family: Orbiliaceae |
| | | Gibberella |
| | | Family: Nectriaceae |
| | | Order: Hypocreales |
| | | Order: Sordariales |
| | | Unidentified |
| | | Sistotrema |
| | | Order: Tremellodendropsidales |
| | | Guehomyces |
| | | Unidentified |
| | | Family: Mortierellaceae |
| | | Unidentified |
| | D3 | Alternaria |
| | | Order: Pleosporales |
| | | Unidentified |
| | | Order: Chaetothyriales |
| | | Aspergillus |
| | | Trichoderma |
| | | Gibberella |
| | | Family: Nectriaceae |
| | | Order: Hypocreales |
| | | Order: Microascales |
| | | Apodus |

| | | |
|---|---|---|
| **Control** | C5 | Order: Hypocreales |
| | | Unidentified |
| | | Schizophyllum |
| | | Order: Tremellales |
| | | Wallemia |
| | | Olpidium |
| | C6 | Alternaria |
| | | Order: Pleosporales |
| | | Unidentified |
| | | Penicillium |
| | | Family: Aspergillaceae |
| | | Oidiodendron |
| | | Verticillium |
| | | Fusarium |
| | | Family: Nectriaceae |
| | | Order: Hypocreales |
| | | Order: Microascales |
| | | Order: Auriculariales |
| | | Guehomyces |
| | | Wallemia |
| | C7 | Alternaria |
| | | Order: Pleosporales |
| | | Penicillium |
| | | Family: Aspergillaceae |
| | | Talaromyces |
| | | Family: Trichocomaceae |
| | | Peziza |
| | | Clonostachys |
| | | Fusarium |
| | | Family: Nectriaceae |
| | | Order: Hypocreales |
| | | Chaetomium |
| | | Humicola |
| | | Zopfiella |
| | | Family: Chaetomiaceae |
| | | Order: Sordariales |
| | | Unidentified |
| | | Family: Ceratobasidiaceae |
| | | Order: Tremellales |
| | | Wallemia |
| | | Spizellomyces |
| | | Claroideoglomus |
| | | Mortierella |
| | | Olpidium |
| | | Unidentified |
| | C8 | Unidentified |
| | | Penicillium |
| | | Family: Aspergillaceae |
| | | Talaromyces |
| | | Pleuroascus |
| | | Peziza |
| | | Fusarium |
| | | Gibberella |
| | | Order: Hypocreales |
| | | Order: Microascales |
| | | Trichocladium |
| | | Family: Lasiosphaeriaceae |
| | | Order: Sordariales |
| | | Unidentified |

| | | |
|---|---|---|
| **Nutrient Addition** | N4 | Order: Auriculariales |
| | | Sistotrema |
| | | Hymenochaete |
| | | Ischnoderma |
| | | Order: Tremellodendropsidales |
| | | Udeniozyma |
| | | Guehomyces |
| | | Itersonilia |
| | | Naganishia |
| | | Order: Tremellales |
| | | Wallemia |
| | | Unidentified |
| | | Mortierella |
| | | Family: Mortierellaceae |
| | N5 | Alternaria |
| | | Order: Pleosporales |
| | | Unidentified |
| | | Exophiala |
| | | Order: Chaetothyriales |
| | | Family: Aspergillaceae |
| | | Family: Trichocomaceae |
| | | Glarea |
| | | Order: Helotiales |
| | | Family: Pyronemataceae |
| | | Meyerozyma |
| | | Fusarium |
| | | Gibberella |
| | | Family: Nectriaceae |
| | | Order: Hypocreales |
| | | Humicola |
| | | Unidentified |
| | | Fibulochlamys |
| | | Sistotrema |
| | | Guehomyces |
| | | Naganishia |
| | | Wallemia |
| | | Unidentified |
| | | Unidentified |
| | | Mortierella |
| | N6 | Alternaria |
| | | Order: Pleosporales |
| | | Unidentified |
| | | Order: Helotiales |
| | | Family: Pseudeurotiaceae |
| | | Family: Pyronemataceae |
| | | Cyberlindnera |
| | | Verticillium |
| | | Fusarium |
| | | Gibberella |
| | | Family: Nectriaceae |
| | | Order: Hypocreales |
| | | Chaetomium |
| | | Humicola |
| | | Family: Chaetomiaceae |
| | | Apodus |
| | | Family: Lasiosphaeriaceae |
| | | Unidentified |
| | | Sistotrema |
| | | Guehomyces |

| | | |
|---|---|---|
| **Water Withholding** | D3 | Order: Sordariales |
| | | Unidentified |
| | | Marasmius |
| | | Cyathus |
| | | Sistotrema |
| | | Guehomyces |
| | | Order: Tremellales |
| | | Unidentified |
| | | Mortierella |
| | D4 | Alternaria |
| | | Order: Pleosporales |
| | | Family: Aspergillaceae |
| | | Verticillium |
| | | Gibberella |
| | | Family: Nectriaceae |
| | | Order: Hypocreales |
| | | Chaetomium |
| | | Conocybe |
| | | Fibulochlamys |
| | | Cyathus |
| | | Order: Agaricales |
| | | Ischnoderma |
| | | Filobasidium |
| | | Wallemia |
| | | Mortierella |
| | | Unidentified |
| | D5 | Penicillium |
| | | Family: Aspergillaceae |
| | | Chaetomium |
| | | Cercophora |
| | | Guehomyces |
| | | Wallemia |
| | | Unidentified |
| | | Mortierella |
| | | Unidentified |
| | D6 | Alternaria |
| | | Order: Pleosporales |
| | | Unidentified |
| | | Family: Aspergillaceae |
| | | Peziza |
| | | Fusarium |
| | | Family: Nectriaceae |
| | | Order: Hypocreales |
| | | Humicola |
| | | Podospora |
| | | Unidentified |
| | | Conocybe |
| | | Parasola |
| | | Guehomyces |
| | | Wallemia |
| | | Mortierella |
| | D7 | Pyrenochaeta |
| | | Alternaria |
| | | Order: Pleosporales |
| | | Penicillium |
| | | Family: Trichocomaceae |
| | | Oidiodendron |
| | | Unidentified |
| | | Meyerozyma |

| | | | | | | | | |
|---|---|---|---|---|---|---|---|---|
| **Control** | C8 | Fibulochlamys | **Nutrient Addition** | N6 | Unidentified | **Water Withholding** | D7 | Acremonium |
| | | Sistotrema | | | Wallemia | | | Fusarium |
| | | Guehomyces | | | Unidentified | | | Family: Nectriaceae |
| | | Itersonilia | | | Ramicandelaber | | | Order: Hypocreales |
| | | Order: Tremellales | | | Mortierella | | | Trichocladium |
| | | Wallemia | | | Olpidium | | | Zopfiella |
| | | Claroideoglomus | | N7 | Alternaria | | | Family: Chaetomiaceae |
| | | Rhizophagus | | | Order: Pleosporales | | | Order: Sordariales |
| | | Order: Paraglomerales | | | Exophiala | | | Unidentified |
| | | Mortierella | | | Penicillium | | | Unidentified |
| | | Unidentified | | | Talaromyces | | | Fibulochlamys |
| | | | | | Botrytis | | | Guehomyces |
| | | | | | Peziza | | | Wallemia |
| | | | | | Chaetosphaeria | | | Spizellomyces |
| | | | | | Clonostachys | | | Funneliformis |
| | | | | | Trichoderma | | | Unidentified |
| | | | | | Fusarium | | | Mortierella |
| | | | | | Gibberella | | | Olpidium |
| | | | | | Family: Nectriaceae | | | |
| | | | | | Order: Hypocreales | | | |
| | | | | | Family: Halosphaeriaceae | | | |
| | | | | | Zopfiella | | | |
| | | | | | Family: Chaetomiaceae | | | |
| | | | | | Order: Sordariales | | | |
| | | | | | Unidentified | | | |
| | | | | | Fibulochlamys | | | |
| | | | | | Order: Agaricales | | | |
| | | | | | Solicoccozyma | | | |
| | | | | | Wallemia | | | |
| | | | | | Spizellomyces | | | |
| | | | | | Unidentified | | | |
| | | | | | Claroideoglomus | | | |
| | | | | | Family: Claroideoglomeraceae | | | |
| | | | | | Family: Glomeraceae | | | |
| | | | | | Unidentified | | | |
| | | | | | Mortierella | | | |

**Table S7: Shared bacterial taxa across treatments (occupancy of 1)**

| OTU.ID | Occ | Domain | Phylum | Class | Order | Family | Genus | Species |
|---|---|---|---|---|---|---|---|---|
| **Shared taxa between control and water withholding-treated seed samples** | | | | | | | | |
| EU231617.1.1548 | 1 | Bacteria | Firmicutes | Bacilli | Bacillales | Bacillaceae | *Bacillus* | NA |
| HQ318731.1.1489 | 1 | Bacteria | Firmicutes | Bacilli | Bacillales | Bacillaceae | *Bacillus* | *B. flexus* |
| HQ323430.1.1541 | 1 | Bacteria | Firmicutes | Bacilli | Bacillales | Bacillaceae | *Bacillus* | *B. megaterium* |
| KP232906.1.1565 | 1 | Bacteria | Firmicutes | Bacilli | Bacillales | Bacillaceae | *Bacillus* | NA |
| **Shared taxa between control and nutrient-treated seed samples** | | | | | | | | |
| EU231617.1.1548 | 1 | Bacteria | Firmicutes | Bacilli | Bacillales | Bacillaceae | *Bacillus* | NA |
| HQ323430.1.1541 | 1 | Bacteria | Firmicutes | Bacilli | Bacillales | Bacillaceae | *Bacillus* | *B. megaterium* |
| KP232906.1.1565 | 1 | Bacteria | Firmicutes | Bacilli | Bacillales | Bacillaceae | *Bacillus* | NA |

*Note: Occ = occupancy; NA = Not Assigned*

**Table S8: Shared fungal taxa across treatments (occupancy of 1)**

| OTU.ID | Occ | Domain | Phylum | Class | Order | Family | Genus | Species |
|---|---|---|---|---|---|---|---|---|
| **Shared taxa between control and water withholding-treated seed samples** | | | | | | | | |
| OTU_24 | 1 | Fungi | Ascomycota | Eurotiomycetes | Eurotiales | Aspergillaceae | *Aspergillus* | *A. niger* |
| OTU_248 | 1 | Fungi | NA | NA | NA | NA | NA | NA |
| OTU_26 | 1 | Fungi | Ascomycota | Eurotiomycetes | Eurotiales | Aspergillaceae | *Aspergillus* | *A. niger* |
| OTU_27 | 1 | Fungi | Ascomycota | Eurotiomycetes | Eurotiales | Aspergillaceae | *Aspergillus* | NA |
| **Shared taxa between control and nutrient-treated seed samples** | | | | | | | | |
| OTU_151 | 1 | Fungi | Ascomycota | Eurotiomycetes | Eurotiales | Aspergillaceae | *Penicillium* | *P. astrolabium* |
| OTU_19 | 1 | Fungi | Ascomycota | Eurotiomycetes | Eurotiales | Aspergillaceae | *Aspergillus* | *A. niger* |
| OTU_24 | 1 | Fungi | Ascomycota | Eurotiomycetes | Eurotiales | Aspergillaceae | *Aspergillus* | *A. niger* |
| OTU_248 | 1 | Fungi | NA | NA | NA | NA | NA | NA |
| OTU_26 | 1 | Fungi | Ascomycota | Eurotiomycetes | Eurotiales | Aspergillaceae | *Aspergillus* | *A. niger* |
| OTU_27 | 1 | Fungi | Ascomycota | Eurotiomycetes | Eurotiales | Aspergillaceae | *Aspergillus* | NA |

*Note: Occ = occupancy; NA = Not Assigned*

### Supplementary Figures

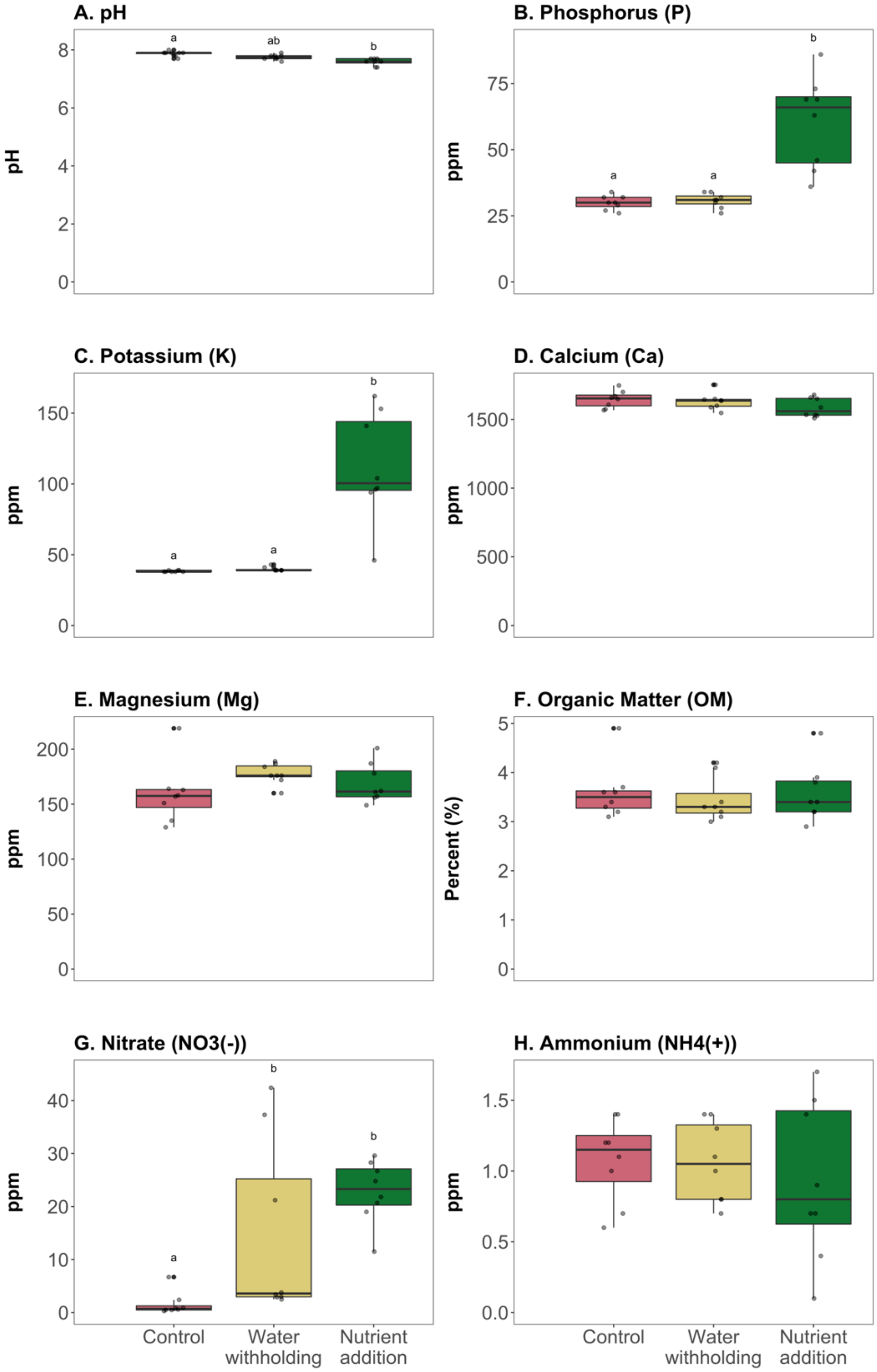


**Figure S1:** Plant rhizosphere soil chemistry for control, water withholding, and nutrient addition treatments of common bean. For each box plot, circles represent one rhizosphere measurement within a treatment. The central horizontal lines represent the mean of measurement, the outer horizontal lines of the box represent the 25th and 75th percentiles. Boxes labelled with different letters are identified as significantly different by a Kruskal-Wallis and post-hoc Dunn's test with a Benjamini-Hochberg false discovery rate correction (p-value significance ranges from <0.05 to <0.0001).

**Supplementary Methods**

**Soil testing at the Michigan State University Soil Plant and Nutrient Laboratory**

Soil samples are dried in a forced air oven at 35°C, then ground with a flail grinder and sieved through a 10 mesh screen.

**pH Determination:** 5 grams of the dried and sieved soil sample is combined with 10 mL distilled water. The mixture is stirred and allowed to stand for 15 minutes. After 15 minutes the mixture is stirred again and immediately tested with a digital pH meter standardized with pH 4.0 and 7.0 buffers.

**Bray P1 Method for Determining Extractable Phosphorous:** 2 grams of dried and sieved soil is combined with 20 mL prepared Bray P1 Working Solution A (0.03 N $NH_4F$ – .025 N HCl) and shaken for 5 minutes at 180-200 rpm. Soil particles are then filtered out of the solution using a Whatman #1 filter. 2 mL of filtered soil extract is then combined with 18 mL prepared Working Solution B (Acid Molybdate – Ascorbic Acid solution) and allowed to sit for 15 minutes while color develops. Color is then read on a spectrophotometer at 660 nm and compared to a standard phosphorous curve to determine concentration.

**Extractable Potassium, Calcium, and Magnesium:** 2.5 grams of prepared soil is combined with 20 mL prepared 20 ml 1N neutral Ammonium Acetate ($NH_4OAc$) extracting solution and shaken for 5 minutes at 180-200 rpm. Soil particles are then filtered out of the solution using a Whatman #1 filter. Nutrient levels are determined using a colorimetric test in a Technicon Autoanalyzer. For magnesium, Magnesium Blue working solution will react with magnesium in the sample solution to develop a blue color. Intensity of the blue color is an indication of the Mg present. For calcium and potassium, sample solutions are mixed automatically with $LiNO_3$ working solution and fed into a flame photometer. Lithium serves as in internal reference standard for determination of Ca and K.

**Organic matter:** 1 gram of prepared soil is combined with 10 ml of $Na_2Cr_2O_7$ solution. 10 ml of 96% concentrated sulfuric acid is added and the mixture is allowed to react for 30 minutes. Mixture is then diluted with 15 mL distilled water and allowed to stand overnight. After standing period, 5 mL supernatant is combined with 5 mL distilled water in a colorimeter tube. Development of orange color in sample is read on a colorimeter calibrated by a standard curve for % organic matter.

**Nitrate Nitrogen:** 10 g dry soil and is combined with 50mL 1 N KCl, shaken for 30 minutes at 180-200 rpm and filtered through a Whatman #2 filter. Nitrate-N content of the filtered soil extract is then determined using either the nitrate-reduction method (QuikChem Method No. 10-107-04-1-A) through a LaChat Rapid Flow Injection Unit, or using a nitrate selective ion electrode connected to an appropriate meter, standardized with prepared nitrate standards.

**Total Nitrogen - Micro-Kjeldahl Block Digestion:** 0.5 g prepared soil is placed in a 100 mL digestion tube with 1 KC-C5 catalyst tablet ( 5 g K2 SO4 + 150 mg CuSO4) and 7 mL concentrated sulfuric acid. Sample is mixed carefully and placed in a block digestor for 1 1⁄2 hours at 375° C. Sample is then removed from digestor block, allowed to cool for 20 to 30 minutes, and filled half full with warm distilled water, thoroughly mixing water with sample. The sample is then filtered through a #2 Whatman filter and analyzed with the LaChat Rapid Flow Injection Unit using the ammonia-salicylate method and appropriate standard.
